# Inhibition of Bruton’s tyrosine kinase reduces NF-kB and NLRP3 inflammasome activity preventing insulin resistance and microvascular disease

**DOI:** 10.1101/745943

**Authors:** Gareth S. D. Purvis, Massimo Collino, Haidee M. A. Tavio, Fausto Chiazza, Caroline E. O’Riodan, Lynda Zeboudj, Nick Guisot, Peter Bunyard, David R. Greaves, Christoph Thiemermann

## Abstract

Activation of inflammatory pathways in myeloid cells initiates insulin resistance leading to the development of type-2 diabetes and microvascular disease. Currently, there are no therapies available that target inflammation in T2D or microvascular disease. In the present study we investigate if Bruton’s tyrosine kinase (BTK) may represent a novel therapeutic target using the FDA approved medication ibrutinib. Ibrutinib treatment protected high fat diet (HFD)-fed mice from developing insulin resistance and improved glycaemic control by restoring signalling through IRS-1/Akt/GSK-3β pathway. These improvements were independent of body weight and calorific intake. Treatment with ibrutinib to mice fed a HFD reduced NF-κB and reduced inflammatory gene expression, this was coupled with decreased activation of the NLRP3 inflammasome in the diabetic liver and kidney. Ibrutinib treatment also protected mice from the development of diabetic nephropathy by reducing monocyte/macrophage infiltration due to reduced expression of the pro-inflammatory chemokines. Ibrutinib treatment to human monocyte derived macrophages significantly reduced pro-inflammatory gene expression and a significant reduction in IL-1β and TNFα after LPS stimulation. In the present study we provide ‘proof of concept’ evidence that BTK is a novel therapeutic target for the treatment of T2D and ibrutinib may be a candidate for drug repurposing in T2D.

## Introduction

The worldwide prevalence of obesity has doubled since 1980 with over 1.9 billion people being overweight or obese (1). The detrimental effects of diet induced obesity culminate in a state of chronic inflammation (2). The control of energy and metabolism in tissues is directed largely by innate immune cells, including macrophages, for the production of soluble effector molecules including cytokines and chemokines (3). This state of ‘low-grade’ chronic inflammation predisposes individuals to metabolic syndrome and the associated comorbidities that develop over time, including, type 2 diabetes (T2D), cardiovascular disease and non-alcoholic fatty liver disease (4)(5). These complications of obesity are a major burden financially on health care providers. The diabetic milieu, is characterised by hyperglycemia, hyperlipidemia, reactive oxygen species, advanced glycation that activate Toll like receptor’s (TLR) in the liver, adipose and on myeloid cells (6). Increased TLR2/4 activation is observed in recently diagnosed type-2 diabetic subjects (7). Downstream signaling through MyD88 activates PKC and MAPK and ultimately leads to the up-regulation of pro-inflammatory pathways via NF-kB (8). Activation of PKC, JNK, and IKKαβ have all been shown to contribute to increased serine phosphorylation on IRS-1 (9)(10)(11), a molecular marker of insulin resistance (12)(13)(14). The resulting hyperglycemia leads to microvascular dysfunction culminating in clinical disease predominantly in the retina and kidney (15)(16).

Myeloid cell activation has been proven to be critical to the development of insulin resistance; ablation of CD11c positive cells (monocytes/macrophages) normalizes insulin sensitivity in obese mice (17). Activation of the NF-κB pathway and the NLRP3 inflammasome pathways have been heavily implicated in orchestrating the inflammatory response in T2D (18)(19). Pharmacological inhibition or genetic deletion of components of the NF-κB pathway i) prevent the development of high fat diet induced insulin resistance (20)(21) and ii) slows the progression of microvascular disease. It has a been demonstrated that myeloid specific deletion of IKK-β, but not deletion in hepatocytes or adipocytes that contributes to the development of insulin resistance (18). Currently there are no medicines available for diabetes that target inflammation and/or prevent the development of microvascular complications. Repurposing existing FDA approved medications that have potent anti-inflammatory effects for new indications could be a cost-effective approach to prevent and/or treat the development of microvascular complications of diabetes. Anti-inflammatory agents have been shown to be powerful tools in pre-clinical models. However, to date little translational research as followed up on these successes.

Bruton’s tyrosine kinase (BTK) is a tyrosine kinase that plays an essential role in B lymphocyte development. Mutations in the BTK gene are implicated in the primary immunodeficiency disease X-linked agammaglobulinemia. As well as, being highly expressed in B-lymphocytes, BTK is also highly expressed in monocytes/macrophages (22), the latter is the key cell types that drives the development of insulin resistance. Importantly, BTK is not expressed in hepatocytes or adipocytes the main cell types effected by obesity induced insulin resistance (23). In monocyte/macrophages BTK has a proposed role in signal transduction downstream of numerous TLR’s. XID mice, which have non-signaling kinase domain in BTK, have reduced activation of PI3K and Atk, upon TLR activation (24)(25)(26). Macrophages from these mice fail to polarize to M1 and have reduced activation to NF-kB signaling following LPS stimulation (27).

The first BTK inhibitor to be FDA approved for the clinical use was ibrutinib for the treatment of chronic lymphatic leukemia (CLL). In pre-clinical models of pro-inflammatory diseases inhibition of BTK has been shown to have therapeutic utility limiting excessive inflammation. BTK inhibition using LMF-A13 improved clinical scores in a murine model of collagen induced arthritis by reducing NF-kB dependent pro-inflammatory cytokine production from B-lymphocytes (28). Ibrutinib treatment was shown to reduce lesion size in a model of murine cerebral ischemia reperfusion injury by reducing IL-1β levels, via reduced NLRP3 inflammasome formation (29).

In the present study we sought to investigate whether the FDA approved medicine ibrutinib could inhibit limit inflammation in a murine model of high fat diet induced T2D, and hence reduce the development of insulin resistance and microvascular disease. We report for the first time that therapeutic intervention with ibrutinib significantly improved glycemic control by restoring normal insulin signalling in diabetic mice. Treatment with ibrutinib protected mice from developing diabetic nephropathy and hepatosteatosis by reducing monocyte/macrophage infiltration and inflammatory gene expression. We demonstrate that inhibition of BTK with ibrutinib attenuates NF-κB dependent gene expression and NLRP3 inflammasome activity in primary murine and human macrophages. Therefore, we propose that ibrutinib may represent a novel drug repurposing candidate in diabetes.

## Results

### Treatment with ibrutinib improves diabetic phenotype in a murine model of insulin resistance

C57BL/6J mice were fed either a chow or HFD for 6 weeks and then treated with ibrutinib (3 or 30 mg/kg; p.o.; 5 times per week) or vehicle for a further 6 weeks. Mice fed a high fat diet gained significantly more weight than mice fed a chow diet, specifically they more gained fat mass (Table 1). Treatment with ibrutinib (3 or 30 mg/kg) to HFD-fed mice did not alter calorific intake, weight gain or cause hepatocellular damage (Table 1). When compared to mice fed a chow diet, mice fed a HFD exhibited a significant elevation in fasted blood glucose (8.62 vs. 12.82 mmol). Mice fed a HFD and treated with either 3 or 30 mg/kg ibrutinib had significantly lower in non-fasted blood glucose compared to mice fed a HFD and treated with vehicle (12.82 vs. 8.04 and 7.71 mmol, respectively) Figure 1A. When compared to mice fed a chow diet, mice fed a HFD exhibited a significant and prolonged increase in serum glucose levels after challenge with oral glucose in the OGTT, (Figure 1A/B), elevated plasma insulin levels (Figure 1C) and elevated non-fasted blood glucose (Figure 1D) indicating the development of diet induced insulin resistance. Mice fed a HFD and treated with either 3 or 30 mg/kg ibrutinib display no change in OGTT when compared to chow fed mice (Figure 1A/B); and significantly lower plasma insulin levels (Figure 1C) and non-fasted blood glucose (Figure 1D) and when compared to mice fed a HFD treated with vehicle. Thus, the BTK inhibitor ibrutinib protects mice from the development of HFD induced insulin resistance independent of altering diet induced obesity.

**Figure 1:**
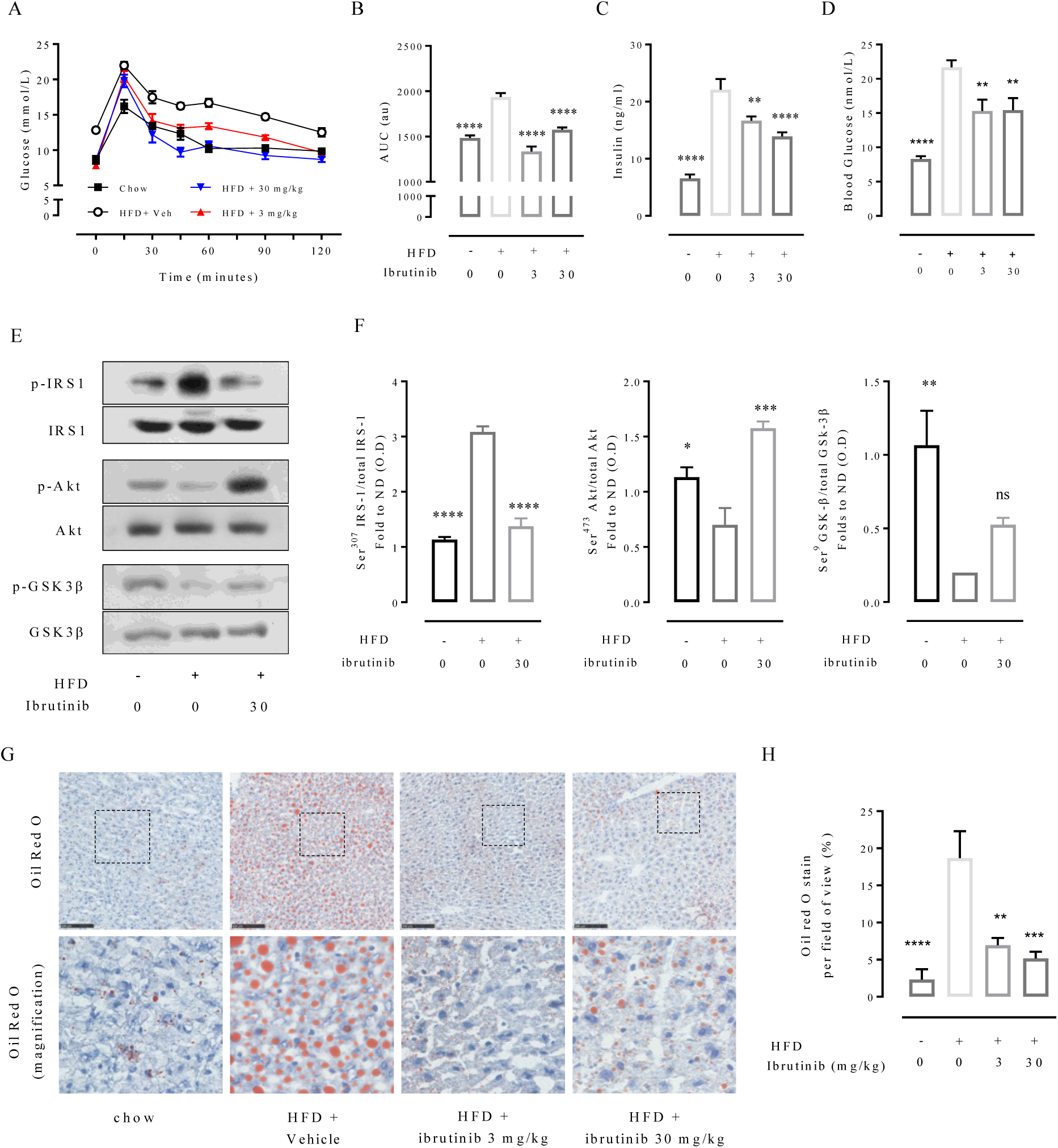
Ibrutinib treatment attenuates the development of HFD induced insulin resistance in mice. C57BL/6 mice, fed a standard diet (chow) or a high-fat diet (HFD) for 12 weeks, were treated with vehicle or ibrutinib (3 or 30 mg/kg) five times per week between weeks 7 and 12. A) Oral glucose tolerance was assessed over 120 min 1 week prior to harvest. B) The area under curve (AUC) of OGTT was calculated for respective groups and used for statistical analysis. C) Serum insulin levels were measured in plasma isolated from whole blood at harvest. D) Basal non-fasted blood glucose was measured at week 12 one hour prior to harvest. E) Representative western blots for phosphorylation of Ser^307^ on IRS-1 in the liver and normalized to total IRS-1; for phosphorylation of Ser^473^ on Akt in the liver and normalized to total Akt; for phosphorylation of Ser^9^ on GSK-3β in the liver and normalized to total GSK-3β and F) quantified using densitometry. G) Representative images of hepatic lipid deposition assessed by Oil Red-O staining and H) quantified. Scale bars measure 50 µm. Data analyses by a one-way ANOVA followed by a Bonferroni *post-hoc* test and the mean is expressed mean± SEM of n=9/10 per group.**p< 0.01, ***p< 0.001, ****p< 0.0001 vs. HFD.

**Table 1:** Basic metabolic parameters measured in interventional study.

|  | **chow** | **HFD +**<br>**Vehicle** | **HFD +**<br>**ibrutinib**<br>**(3 mg/kg)** | **HFD +**<br>**ibrutinib**<br>**(30 mg/kg)** |
| --- | --- | --- | --- | --- |
| **Body weight**<br>**(g)** | 31.24±0.4316* | 36.1±0.8 | 33.55± 0.7863 | 31.21±0.3078 |
| **Weight gain**<br>**from baseline (g)** | 7.0 ±0.5* | 11.2±0.5 | 9.4±0.4 | 9.9±0.8 |
| **Food intake per**<br>**week (g)** | 29.16±0.93 | 18.14±0.67 | 16.83±0.92 | 16.674±0.89 |
| **Calorific intake**<br>**per week (kCal)** | 105.22±4.04 | 100.85±3.39 | 93.62±4.60 | 93.18±4.32 |
| **Kidney as % of**<br>**body weight** | 1.12±0.02 | 0.91±0.02 | 0.89±0.02 | 0.94±0.03 |
| **Liver weight as**<br>**% of body**<br>**weight** | 4.45±0.20* | 3.23±0.07 | 3.54±0.08 | 4.32±0.04 |
| **Inguinal fat as**<br>**% of body**<br>**weight** | 0.45±0.073* | 1.30±0.093 | 1.17±0.069 | 1.20±0.12 |
| **Epididimal fat as**<br>**% of body**<br>**weight** | 2.46±0.27* | 4.98±0.35 | 4.30±0.32 | 4.08±0.19 |
| **ALT**<br>**(U/L)** | 17.99±1.792* | 61.04±13.16 | 58.47±9.031 | 54.63±7.16 |
| **AST**<br>**(U/L)** | 80.88±5.88* | 120.4±16.55 | 115±11.87 | 119±12.18 |

Body weight (g), body weight gain from baseline (g), Food intake per week (g), caloric intake per week, inguinal fat weight, epididymal fat weight, ALT and AST were measured in mice fed a standard diet (chow) or a high-fat diet (HFD) for 12 weeks. HFD-mice were treated with either vehicle or ibrutinib (3 or 30 mg/kg p.p) five times per weeks between weeks 7 and 12. Data are expressed as mean± SEM of 9/10 mice per group. Data were analyzed by a one-way ANOVA followed by a Bonferroni post-hoc test, \*p < 0.05 vs. WT + HFD.

We next investigated alterations in the insulin signaling pathway caused by HFD. When compared to mice fed a chow diet, mice fed a HFD treated with vehicle have an increase in the phosphorylation on serine 307 of IRS-1 (Figure 1E/F), resulting in a reduction in phosphorylation of downstream mediators Akt on serine 473 and GSK-3β on serine 9 (Figure 1E/F). Mice fed a HFD and treated with ibrutinib demonstrated less phosphorylation of serine 307 of IRS-1, attenuating the decrease in phosphorylation of Akt and GSK-3β compared to mice fed a HFD (Figure 1E/F). Thus, ibrutinib treatment preserved insulin signaling through IRS1 in mice subjected to a HFD resulting in a higher degree of activation of the Akt survival pathway.

The development of peripheral insulin resistance is associated with the development of hepatosteatosis. We therefore quantified fat deposition in the liver using oil-red O staining. When compared to chow diet, mice fed a HFD (treated with vehicle) had a significant increase in oil red O staining in the liver (Figure 1G/H). Treatment of HFD mice with ibrutinib (3 or 30 mg/kg) resulted in a significant reduction in Oil red O staining in the liver (Figure 1G/H), demonstrating less lipid deposition in the liver and hence reduced hepatic steatosis.

### Treatment with ibrutinib reduces inflammation via inhibition of NF-κB and the NLRP3 inflammasome

We next wanted to investigate if the improvements in the diabetic phenotype seen in mice treated with the ibrutinib could be attributed to reduced activation of pro-inflammatory pathways. When compared to mice fed a chow diet, liver tissues from mice fed a HFD exhibited a significant increase in the phosphorylation of Ser^32/36^ on IκBα and, hence, activation (phosphorylation) of the IKK complex, allowing for increased nuclear translocation of the NF-κB subunit p65 to the nucleus where its acts a transcription factor (Figure 2A/B). Indeed, we observed increases in the expression of pro-inflammatory cytokines (IL-1β, IL-18, TNFα and IL-10) (Figure 2C). Liver tissues from mice fed a HFD and treated with ibrutinib showed a significant reduction in the degree of phosphorylation of IκBα on Ser^32/36^ and, hence, reduced activation of the IKK complex and as a consequence reduced nuclear translocation of the p65 NF-κB subunit to the nucleus (Figure 2A/B). As a result, ibrutinib treatment to mice fed a HFD attenuated the increase in the gene expression of pro-inflammatory cytokines (IL-1β, IL-18, TNFα and IL-10) in the liver (Figure 2C).

**Figure 2:**
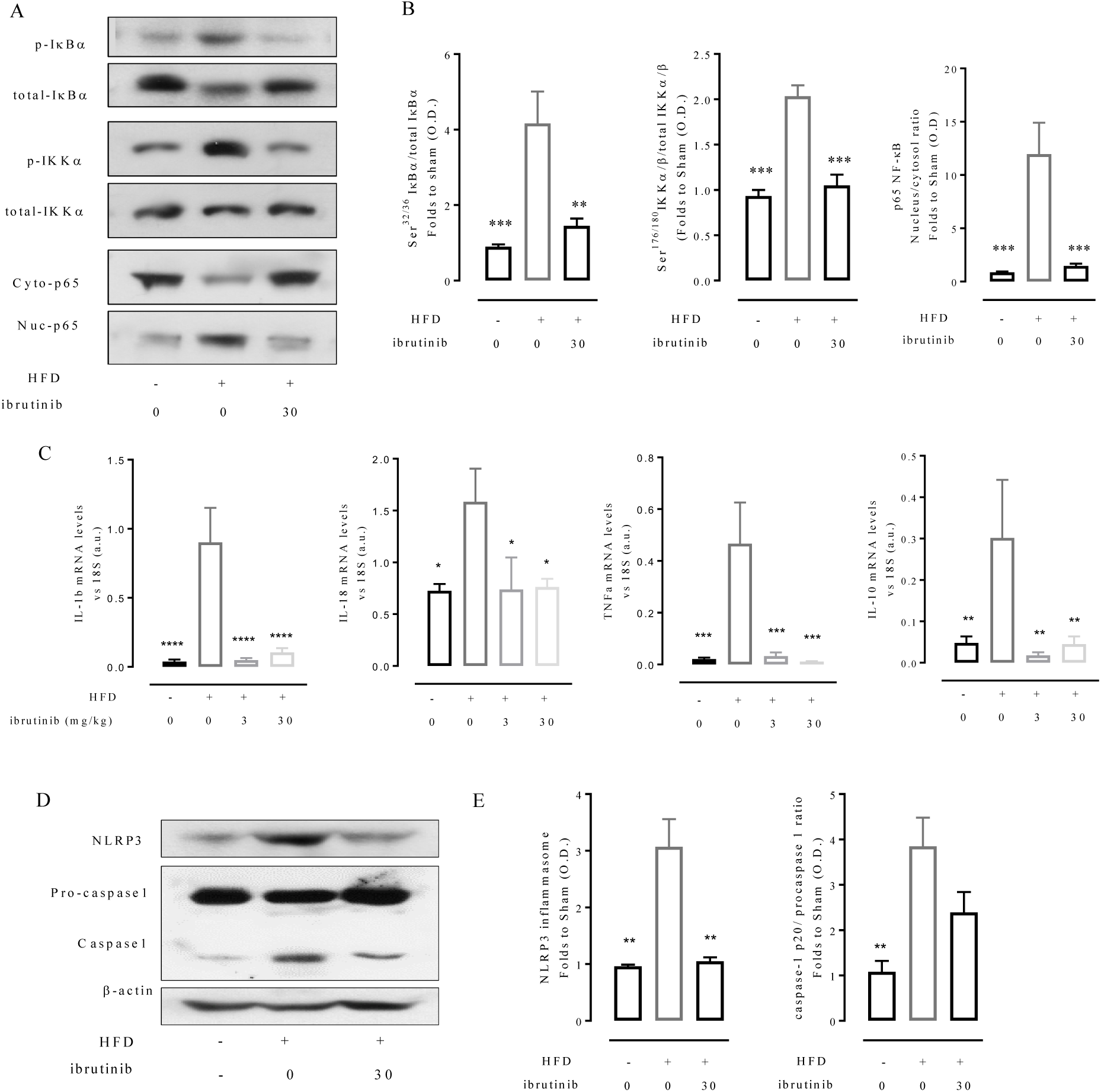
Ibrutinib treatment reduces inflammation in the diabetic liver via inhibition of NF-κB and the NLRP3 inflammasome. C57BL/6 mice, fed a standard diet (chow) or a high-fat diet (HFD) for 12 weeks, were treated with vehicle or ibrutinib (3 or 30 mg/kg) five times per week between weeks 7 and 12. A) Representative western blots for phosphorylation of Ser^32/36^ on IKBα in the liver and normalized to total IKBα; for phosphorylation of Ser^176/180^ on IKKα in the liver and normalized to total IKKα; nuclear translocation of p65 and B) quantified using densitometry. C) Relative gene expression of *IL-1β, IL-18, TNFα* and *IL-10* were assessed by qPCR and normalized to *18S*. D) Representative western blots for NLRP3 inflammasome assembly and the proteolytic cleavage of pro-caspase 1 to caspase 1 normalized to β-actin and E) quantified using densitometry. Data analyses by a one-way ANOVA followed by a Bonferroni *post-hoc* test and the mean is expressed mean± SEM of n=6-10 per group. **p< 0.01, **p<0.01, ***p<0.001 and ****p<0.0001.

When compared to mice fed a chow diet, liver tissues from mice fed a HFD demonstrated assembly of the NLRP3 inflammasome, which results in the proteolytic cleavage of pro-caspase 1 into caspase 1 (Figure 2D/E). Mice fed a HFD and treated with ibrutinib have significantly reduced formation of the NLRP3 inflammasome and, hence, reduced proteolytic cleavage of pro-caspase 1 to caspase 1 in the liver (Figure 2D/E). Collectively, these results demonstrate that chronic treatment with ibrutinib reduced NF-κB activation and formation of the NLRP3 inflammasome in the liver, thus reducing pro-inflammatory gene expression in the liver, which are the main drivers of the initiation and development of peripheral insulin resistance.

### Treatment with ibrutinib protects mice from the development of diabetic nephropathy

The pro-inflammatory environment coupled with insulin resistance in T2D leads to the development of microvascular complications overtime including diabetic nephropathy. Mice fed a HFD (treated with vehicle) had an increased urinary albumin to creatinine ratio (ACR), a key pathophysiological read out for proteinuria due to diabetic nephropathy (Figure 3A). This was coupled with classical histological changes in the kidney that are consistent with the development of microvascular damage i.e. glomerular hypertrophy, basement membrane expansion and loss of brush borders at the level of the S1-S2 segment of the proximal convoluted tubules (Figure 3B). Prolonged treatment of HFD-mice with ibrutinib (3 or 30 mg/kg) attenuated both the proteinuria and the associated histological signs of glomerular and tubular degeneration (Figure 3A/B). Infiltration of inflammatory macrophages is a hallmark of diabetic nephropathy. Indeed, mice fed a HFD (treated with vehicle) exhibited an increased number of F4/80^+^ macrophages in the kidney which could be attenuated with ibrutinib treatment (Figure 3C/D). Additionally, ibrutinib treatment also attenuated the activation of both the NF-kB and NLRP3 inflammasome in the diabetic kidney (Sup Figure 1). Mice fed a HFD expressed high levels of innate immune cells chemoattractant (CXCL1, CCL2 and CCL5), which could be attenuated with ibrutinib treatment (Figure 3E). Taken collectively our data demonstrate that oral dosing with ibrutinib protects against the development of diabetic nephropathy by reducing immune cell recruitment and inhibition of pro-inflammatory pathways.

**Figure 3:**
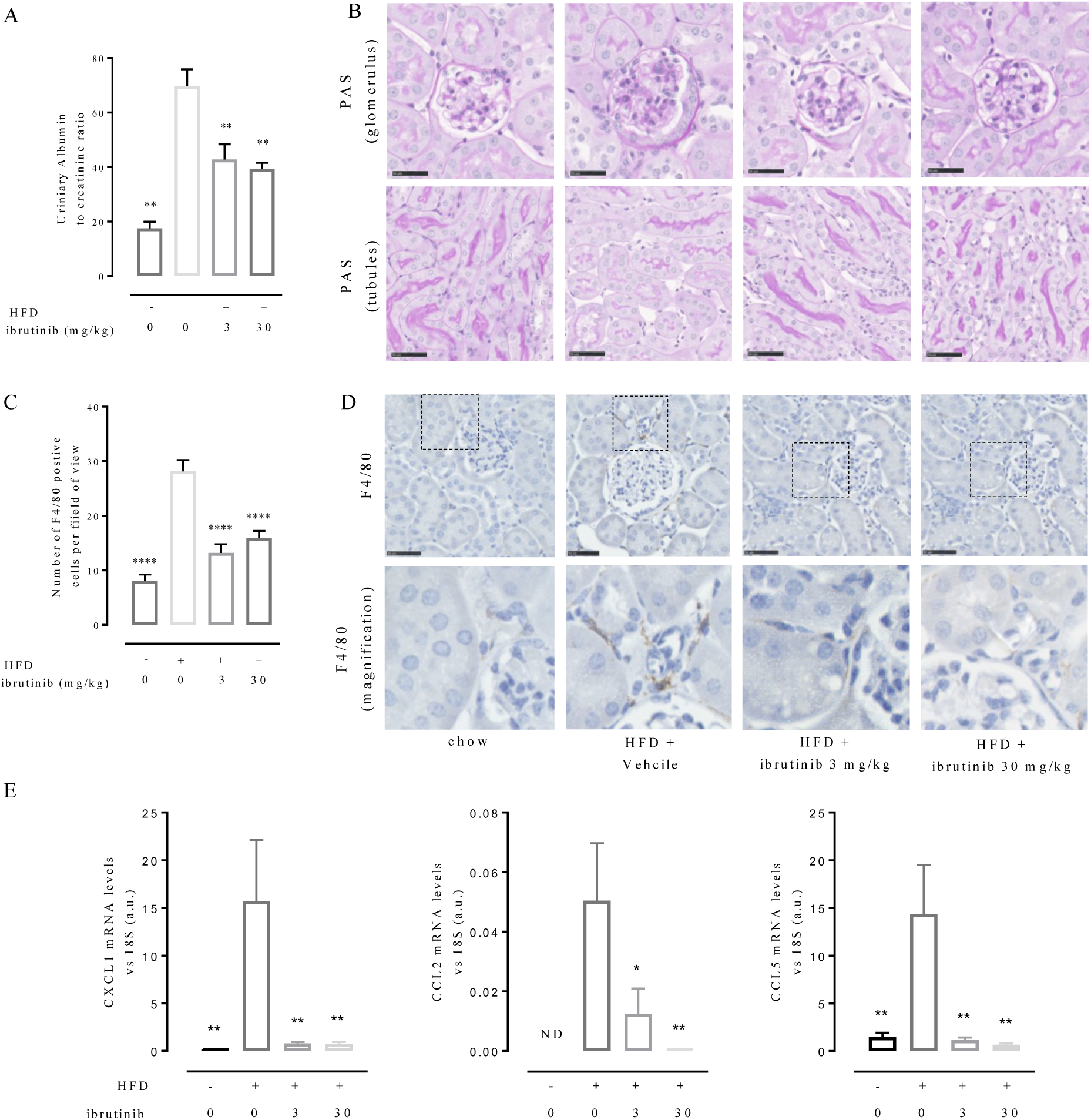
Ibrutinib treatment prevents the development HFD induced diabetic nephropathy. C57BL/6 mice, fed a standard diet (chow) or a high-fat diet (HFD) for 12 weeks, were treated with vehicle or ibrutinib (3 or 30 mg/kg) five times per week between weeks 7 and 12. A) Renal function was assessed by urinary albumin to creatinine ratio (ACR) from 18 h urine samples. B) Kidney sections were stained with Periodic acid-Schiff staining to assess histological changes. D) Macrophage infiltration was assessed by immunohistochemical staining for macrophage marker F4/80. and C) quantified. E) Relative gene expression of *CXCL1, CCL2* and *CCL5* were assessed by qPCR and normalized to *18S*. Scale bars measure 50 µm. Data analyses by a one-way ANOVA followed by a Bonferroni *post-hoc* test and the mean is expressed mean± SEM of n=9/10 per group. *p<0.05, **p< 0.01, ****p< 0.0001 vs. HFD.

### Treatment with ibrutinib impairs leukocyte recruitment during acute inflammation

Having demonstrated that chronic treatment with ibrutinib reduced F4/80^+^ macrophage infiltration into the diabetic kidney, we next wanted to investigate the effect of ibrutinib on leukocyte recruitment *in vivo* in a model of acute resolving inflammation. Male C57BL/6J mice were pre-treated with ibrutinib (10 mg/kg; p.o) or vehicle 1 h before being injected with zymosan (100 ug; i.p.). Peritoneal exudates were harvested at 2, 4, 16 and 48 h post zymosan challenge (Figure 4A and B).

**Figure 4:**
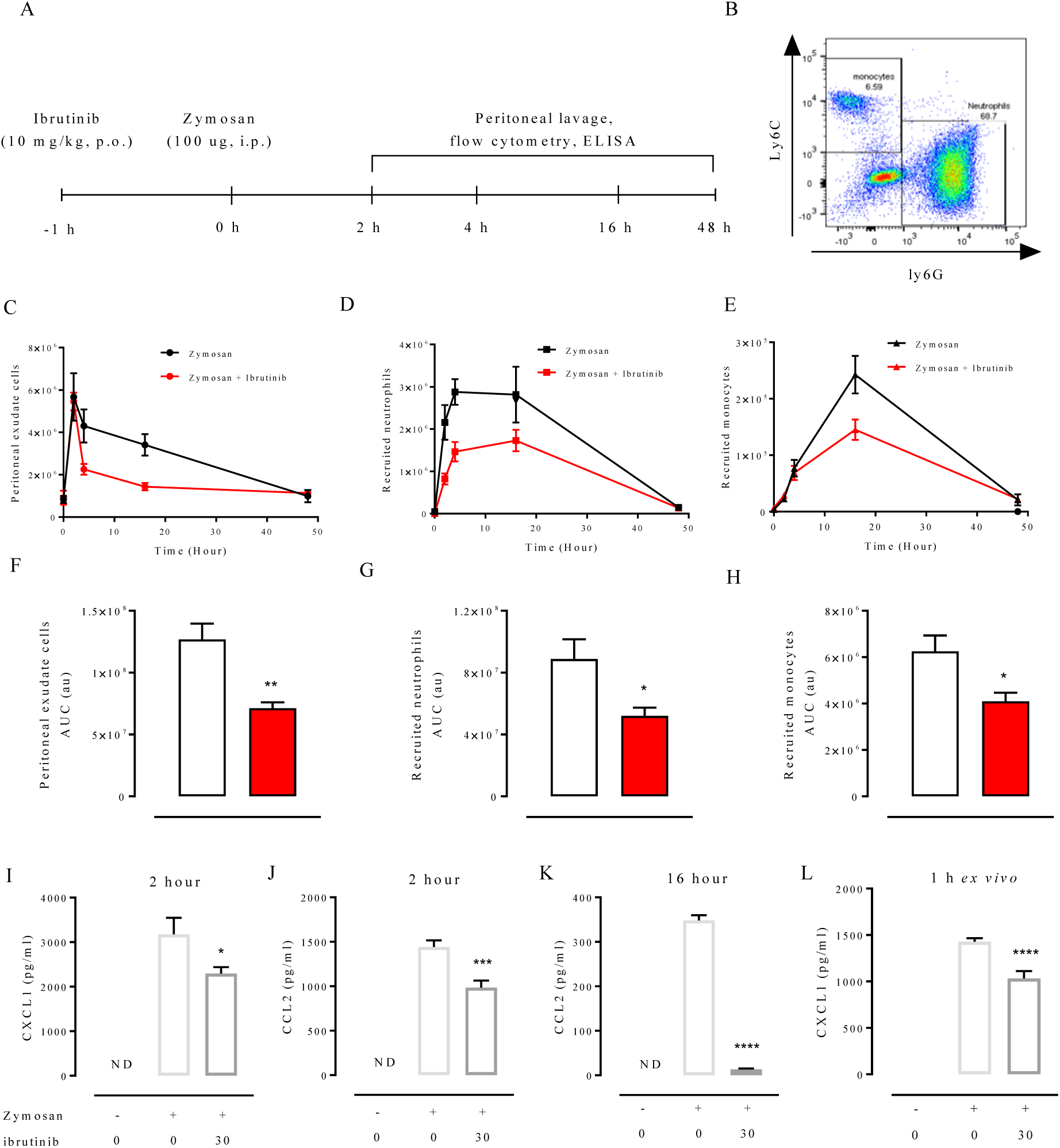
Ibrutinib treatment inhibits inflammatory cells recruitment. C57BL/6 mice were pre-treated with ibrutinib (10 mg/kg, p.o.) or vehicle 1 h prior to zymosan challenge (100 µg, i.p.). Mice were euthanized at 2, 4, 16 and 48 hr and peritoneal lavage samples were collected. A) Diagram of the experimental design. B) Representative flow cytometry plots of neutrophils characterized as CD11b^+^/Ly6C^-^/Ly6G^+^ and monocytes characterized as CD11b^+^/ Ly6C^hi/lo^/Ly6G^-^ from peritoneal lavage fluid. C) Time course of total peritoneal exudate cells and quantified by AUC (F). D) Time course of recruited neutrophils and quantified by AUC (G). E) Time course of recruited monocytes and quantified by AUC (H) White bars: zymosan + vehicle; Red bars: zymosan + ibrutinib. I-K) CXCL1 (at 1 hours post ZIP) and CCL2 (at 2 and 16 h post ZIP) were measured in peritoneal lavage fluid by ELISA at . L) CXCL1 was measured in supernatants from peritoneal macrophages *ex vivo,* after 1 hours of zymosan stimulation ± ibrutinib (10 mM), by ELISA. Data analyses by t-test were there are only two variables or a one-way ANOVA followed by a Bonferroni *post-hoc* test were multiple variables and the mean is expressed mean± SEM of n=6-12 per group. *p<0.05, **p< 0.01, ***p<0.001 and ****p<0.0001.

Compared to mice treated with vehicle, mice treated with ibrutinib and challenged with zymosan had significantly reduced total cell counts, and fewer recruited neutrophils and monocytes over the entire time course (Figure 4C-H). There was a significant reduction in recruited neutrophils at 4 h and 16 h post zymosan in ibrutinib treated mice (Sup Figure 2A/B), and a reduction in recruited monocytes numbers at 16 h (Sup Figure 2D). The chemokines CXCL1 (at 1 h) and CCL2 (at 4 h and 16 h) were measured in peritoneal exudates: CXCL1 was significantly reduced at 2 h and CCL2 levels were lower at 4 hour and 16 h in mice treated with ibrutinib prior to zymosan challenge (Figure 4I-K). To test whether systemic ibrutinib treatment affects the ability of resident peritoneal macrophages to secrete CXCL1, resident peritoneal macrophages were isolated and treated *ex vivo* with ibrutinib prior to zymosan challenge. Secretion of CXCL1 was lower in resident peritoneal macrophages treated with ibrutinib prior to zymosan challenge (Figure 4L). Our data support a model in which ibrutinib reduces the secretion of CXCL1 by resident peritoneal macrophages, which, in turn, reduces neutrophil recruitment leading to less secretion of CCL2, the primary chemoattractant for monocytes.

### Ibrutinib treatment reduces NF-kB and NLRP3 inflammasome activation in murine macrophages

Macrophages are a key cell type involved in the local amplification of the immune response in tissues. Therefore, we wanted to confirm that ibrutinib inhibits the activation of both NF-κB and the inflammasome in macrophages. To investigate this, we first used a macrophage cell line stably transfected with a secreted alkaline phosphatase (SEAP) reporter gene under the transcription regulation of NF-κB and AP1. Macrophages stimulated with LPS for 6 hours exhibited a strong induction of NF-κB (Figure 5A), which could be inhibited when macrophages were pre-incubated with ibrutinib (1 – 30 uM) to inhibit BTK 60 min prior to LPS stimulation (Figure 5A). No cytotoxicity was observed in macrophages treated with up to 30 uM of ibrutinib for 9 hours in an MTT assay (Sup Figure 3A). We then wanted to confirm that BTK inhibition with ibrutinib resulted in less NF-κB activation in primary murine bone marrow derived macrophages (BMDM) (Sup Figure 3B). LPS stimulation induced cytoplasmic to nuclear translocation of p65 in BMDM, which was inhibited by pre-treatment with ibrutinib (10 uM; for one hour prior to LPS stimulation; Figure 5B). Stimulation of BMDMs with LPS also resulted in the up-regulation of pro-inflammatory gene expression including IL-1β, TNFα and IL-6 which was also reduced by pre-treating with ibrutinib, in a dose-dependent manner (Sup figure 3C).

**Figure 5:**
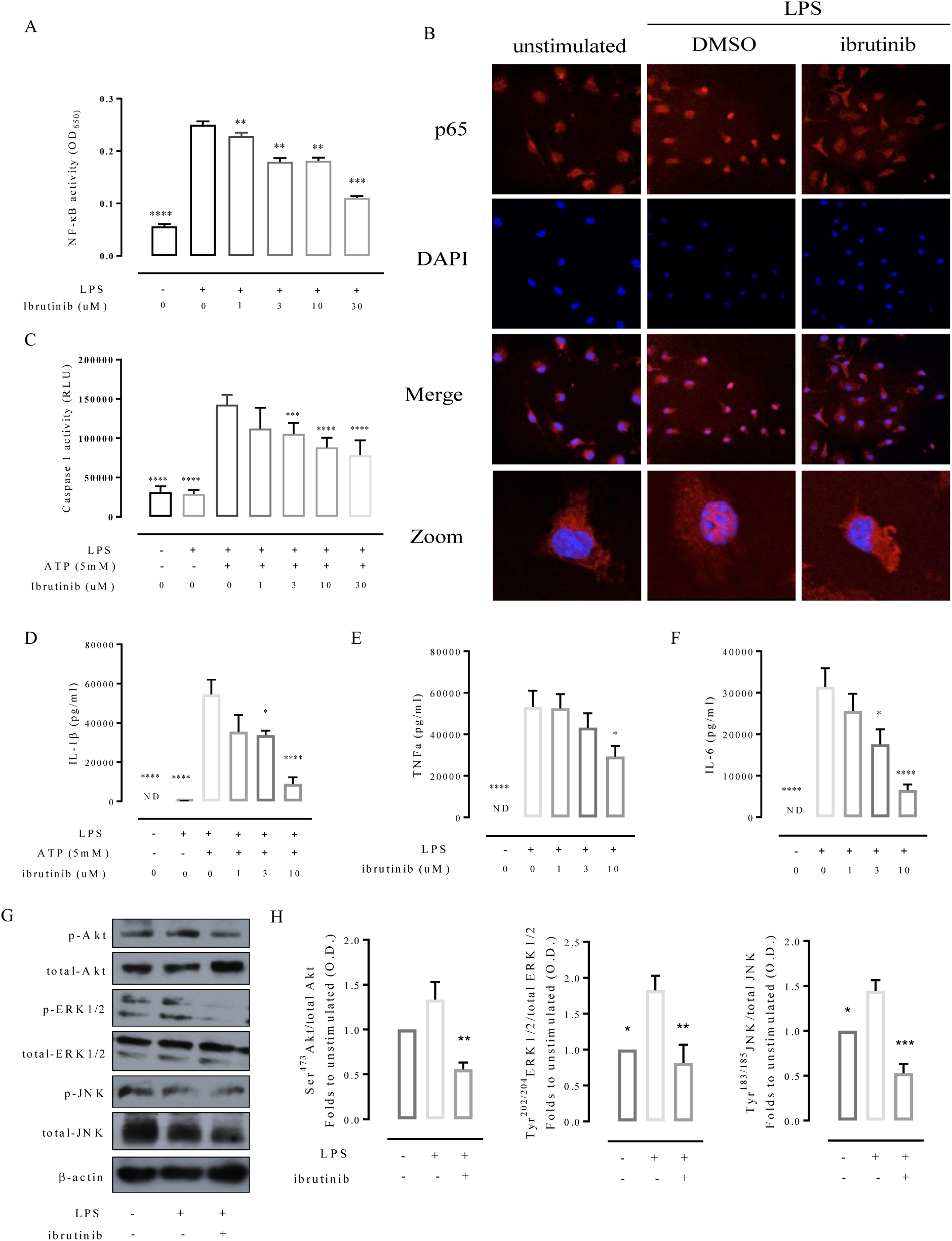
Ibrutinib treatment reduces NF-κB and NLRP3 activation in macrophages. A) Raw Blue cells (macrophage cell line) stably transfect with a secreted alkaline phosphatase (SEAP) under the transcription regulation of NF-KB and AP1 were pre-treated with ibrutinib (1-30 µM) or vehicle (0.6 % DMSO) for 1 hour prior to LPS (0.1 µg ml^-1^) stimulation, after 6 hours SEAP was quantified by at OD_650_. Summary data of n=4 independent experiements. B) Bone marrow derived macrophages (BMDMs) pre-treated with ibrutinib (10 µM) prior to LPS (0.1 µg ml^-1^) stimulation 30 minutes followed by p65 staining. Confocal microscopy images are illustrative of two separate experiments. C) BMDMs were pre-treated with ibrutinib (1-30 µM) or vehicle (0.6 % DMSO) for 1 hour prior to LPS (0.1 µg ml^-1^) stimulation for 8 h (in the last 1 h ATP was added to activate caspase 1); caspase activity was measured using luminescence Summary data of n=4 independent BMDM preparations. D-F) BMDMs pre-treated with ibrutinib (1-30 µM) or vehicle (0.6 % DMSO) for 1 hour prior to LPS (0.1 µg ml^-1^) stimulation for 8 h and cytokine (IL-1β, TNFα and IL-6) secretion measured by ELISA (in the last 1 h ATP was added to IL-1β secretion) Summary data of n=6 independent BMDM preparations. G) Representative western blots for phosphorylation of Ser^473^ on Akt in the BMDM and normalized to total Akt; for phosphorylation of Thr^202^/Tyr^204^on ERK1/2 in the BMDM and normalized to total ERK1/2 for phosphorylation of Thr^183^/Tyr^185^ on JNK in the BMDM and normalized to total JNK; and H) quantified using densitometry. Data analyses by a one-way ANOVA followed by a Bonferroni *post-hoc* test and the mean is expressed mean± SEM. *p<0.05,**p< 0.01, ***p<0.001 and ****p<0.0001.

Next, we investigated if BTK inhibition with ibrutinib could block the formation of the NLRP3 inflammasome in primary murine BMDM using a caspase-1 activity assay. BMDMs treated with LPS to do not activate caspase-1 (Figure 5C), as a second signal is required. To provide this second stimulus, ATP (5 mM) was added to BMDM in the last 60 min of LPS challenge, which resulted in a significant increase in caspase-1 activity (Figure 5C). Pre-treatment of BMDM with ibrutinib (1-30 uM) 60 min prior to LPS + ATP stimulation resulted in a dose-dependent inhibition of caspase-1 activity (Figure 5C).

In order to confirm that the observed changes in macrophage gene expression led to changes in pro-inflammatory cytokine secretion. BMDM were stimulated with LPS for 8 hours and supernatants collected and cytokine levels measured by ELISA; for IL-1β secretion ATP was added in the last 60 mins. Pre-treatment with ibrutinib min prior to LPS stimulation significantly inhibited the secretion of pro-inflammatory cytokines IL-1β, TNFα, IL-6 in a dose-dependent manner (Figure 5D/E/F). To investigate the effect of BTK inhibition on macrophage signaling during the inflammatory response, we focused on protein kinase B (Akt) and mitogen-activated protein kinase (MAPK) activation. BMDM stimulated with LPS for 15 min exhibited an increase the degree of phosphorylation on Ser^473^ of Akt, tyr^204^/thr^202^ of ERK1/2 and thr^183^/tyr^185^ on JNK, which could be inhibited by pre-treatment with ibrutinib (10 uM) for 60 min prior to LPS stimulation (Figure 5G/H). These experiments clearly demonstrate that inhibition of BTK with ibrutinib has potent anti-inflammatory effects, resulting in the inhibition of both NF-κB signaling and NLRP3 inflammasome activity in macrophages.

### Ibrutinib treatment reduces pro-inflammatory gene expression and cytokines secretion in human monocytes derived macrophages

To test the translational relevance of our findings obtained in murine macrophages, we studied the activation of pro-inflammatory gene expression and the secretion of pro-inflammatory cytokines in human monocyte derived macrophages (hMoDM). Human monocytes were isolated from leukocyte cones obtained from healthy volunteers and enriched by column seperation (Sup Figure 4), and then differentiated into macrophages. On day 7 of differentiation, hMoDM were pre-treated with LPS to induce inflammatory gene expression. Pre-treatment with ibrutinib (10 uM) 60 min prior to LPS stimulation inhibited the increase in pro-inflammatory gene expression (IL-1β and TNFα) seen in hMoDM treated with LPS alone (Figure 6A/B). Pre-treatment with ibrutinib prior to LPS stimulation significantly inhibited the secretion of both NF-kB and NLRP3 dependent cytokines TNFα and IL-1β by 10-fold and 2-fold respectively (Figure 6C/D).

**Figure 6:**
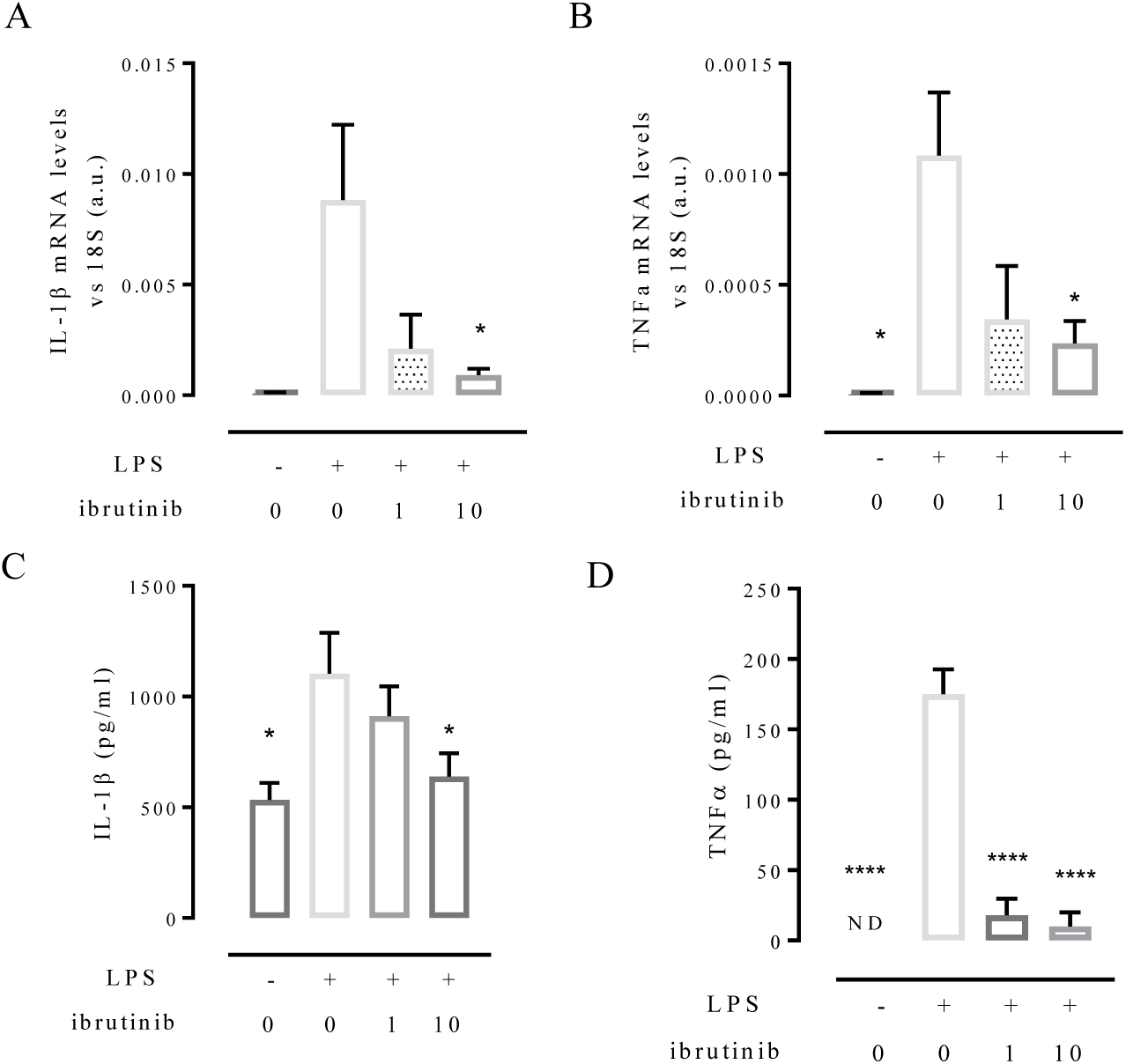
Ibrutinib reduces pro-inflammatory gene expression and cytokine secretion in human macrophages. Human monocytes are isolated from leukocyte cones and purified by magnetic bead purification. A) Representative flow cytometry plots showing monocyte enrichment; monocytes are defined as CD14^+^/CD16. hMoDM are pre-treated with ibrutinib (10 µM) or vehicle (0.6 % DMSO) for 1 hour prior to LPS (0.1 µg ml^-1^) stimulation for 8 h (in the last 1 h ATP was added to IL-1β secretion). B) Relative gene expression of *IL-1β* and*, TNFα* were assessed by qPCR and normalized to 18S. C) Cytokine secretion was assessed by Bead based ELISA kit. Data analyses by a one-way ANOVA followed by a Bonferroni *post-hoc* test and the mean is expressed mean± SEM of n=12 donors. *p<0.05 and ****p<0.0001.

## Discussion

In this study, we set out to investigate if the anti-cancer medicine ibrutinib could represent a novel drug repositioning opportunity for the treatment of diet induced insulin resistance and microvascular disease. The main findings of this in pre-clinical interventional study are that chronic treatment of mice on a HFD with ibrutinib (1) improves insulin sensitivity, lowering blood insulin and blood glucose levels via restoration of IRS-1/Akt/GSK-3β signalling pathway (in the liver), independent of changes in body weight or calorific intake; (2) reduces the activation of both NF-κB and NLPR3 inflammasome in both the kidney and liver, (3) protecting mice from the development of liver hepatosteatosis and diabetic nephropathy. Chronic ibrutinib treatment was associated with a reduction in macrophage accumulation in the kidney of diabetic mice. Mechanistically we demonstrate that systemic inhibition of BTK with ibrutinib reduces immune cell infiltration in sterile peritonitis, by reducing the secretion of the chemoattractants CXCL1 and CCL2 from resident peritoneal macrophages. We demonstrate in primary murine and human macrophages that ibrutinib reduces LPS-stimulated NF-κB activation and attenuates caspase-1 activity, resulting in a reduction in pro-inflammatory gene expression and a reduction in secreted cytokines (IL-1β and TNFα). Critically, our work has identified a novel therapeutic target in macrophage expressed BTK. Importantly, we provide ‘proof-of-concept’ data that an existing, FDA approved drug protects mice fed a HFD from the development of insulin resistance, by reducing inflammation and, hence protecting mice from microvascular damage.

The observed impairment in glucose tolerance seen in mice fed a HFD is, mediated, in part, by alterations in insulin signalling in the liver. The IRS-1/Akt/GSK-3β signalling cascade is a key regulator of glucose transportation, energy metabolism and ultimately tissue protection (30)(31). A marker of peripheral insulin resistance is the phosphorylation of Ser^307^ on IRS-1; this uncouples IRS-1 from the insulin receptor blunting the ability of insulin to signal through its receptor, which, in turn, reduces GLUT4 translocation to the cell surface to facilitate glucose uptake in peripheral organs, leading to hyperglycemia (20)(32). It should be noted that although phosphorylation of Ser^307^ is a widely used marker of insulin resistance, it is not causative *in vivo* (14). When compared to mice fed a chow diet, mice fed a HFD showed increased phosphorylation of Ser^307^ on IRS-1 (in the liver), while treatment of HFD-mice with ibrutinib attenuated this increase in the phosphorylation of Ser^307^ on IRS-1 resulting in restoration of normal blood glucose and insulin levels and an improved OGTT. The subsequent re-activation of the downstream mediator of IRS-1, Akt, has been shown to confer organ protection in many pre-clinical models associated with inflammation including sepsis (33)(34), hemorrhagic shock-induced organ dysfunction (35)(36), and diabetes (20)(37). Indeed re-activating GSK-3β is associated with increased conversion of glucose to glycogen, which could also be a factor contributing to the observed glucose lowering effects *in vivo,* and is a key regulator of NF-κB activation (38)(39). In the present study we demonstrate inhibition of BTK with ibrutinib protects against the development of insulin resistance by restoring insulin signalling, most likely due to reduced inflammation, in myeloid cells.

Myeloid cell activation has been consistently shown to be the driver of the development of insulin resistance through activation of numerous pathways including NF-κB. Interventional studies have reported the beneficial effects NF-κB inhibition on the development of insulin resistance (20)(40)(41). However, in these published studies it is unclear what the cellular target was. Mice with myeloid specific deletion of IKK-β and fed a HFD do not development insulin resistance; while mice fed a HFD with hepatocyte specific deletion of IKK develop less severe insulin resistance (18)(42). Skeletal muscle deletion of NF-kB activity does not protect mice from the development of insulin resistance (43). Taken together these published studies demonstrate that while systemically inhibiting the NF-KB pathways does protect from the development of insulin resistance, it is the myeloid specific inhibition that is the critical cellular target *in vivo*. Here we demonstrate that inhibition of BTK, which has significantly higher expression levels in monocyte/macrophages, compared to the liver, adipose or skeletal muscle (22), results in reduced activation of the NF-kB pathway in both the kidney and liver, thus protecting mice from developing insulin resistance. IKK-β can also regulate insulin sensitivity through direct phosphorylation of IRS-1 suggesting a transcription-independent mechanism of action (18). Indeed, here we show mice fed a HFD and treated with ibrutinib have less activation of IKK-β which correlates with a reduction in IRS-1 phosphorylation, indicative of reduced insulin resistance. We demonstrate for the first time a mechanistic link between BTK, IKK-β and IRS-1 signalling.

To prove the effects of systemic administration of ibrutinib to mice was due to targeting macrophage activation, we demonstrate that in primary murine and human macrophages that ibrutinib treatment, can regulate the inflammatory response at two crucial check-points, both signal one (inhibiting NF-κB dependent transcription of pro-inflammatory genes) and signal two (inhibiting NLRP3 inflammasome activity). Signal one induces the expression of *IL-1β* and *IL-18* mRNA via activation of NF-κB, and signal 2, the activation of NLRP3 inflammasome and the proteolytic cleavage of pro-IL-1β and IL-18 to their active forms. Using the TLR2/4 agonist LPS we demonstrate the BTK inhibition results in less activation of NF-KB, resulting in less nuclear translocation of the p65 sub-unit of NF-κB to the nucleus attenuating the production of pro-inflammatory cytokines and chemokines. BTK has been shown to be involved in the regulation of NF-κB dependent cytokine production in B lymphocytes (44)(45); and BTK uses a positive auto**-**regulatory feedback mechanism to stimulate transcription from its own promoter via NF-κB in B-cells (46). Our data is consistent with work by Gabhann *et al.* who suggest BTK is necessary for TLR4 signal transduction in macrophages (27).

BTK has been proposed to be an essential for the formation of the ASC a key process in the formation of the NLRP3 inflammasome. Ito *et al.* and others have found that the BTK inhibitor LFM-A13 inhibited mature IL-1β secretion from macrophages stimulated with the NLRP3 inflammasome activators (47). In BMDMs we demonstrate that BTK inhibition with ibrutinib prior to LPS stimulation inhibited caspase 1 activity in a dose dependent manner, resulting in less secretion of IL-1β from both murine and human macrophages. Taken together our in vivo and in vitro data demonstrate that ibrutinib targets macrophages reducing the activation of pro-inflammatory pathways.

Even with careful management of blood glucose and strategies to lower circulating lipids microvascular complications develop over time in patients with diabetes. These complications occur predominantly in tissues where glucose uptake is insulin-independent (kidney, retina and the endothelium), as these tissues are exposed to glucose levels close to blood glucose levels. The extent of inflammatory cell accumulation in the diabetic kidney has been associated with the decline of renal function, suggesting a causative link (48). We speculated that BTK is a pivotal kinase in macrophages needed for the production of chemokines that recruitment of immune cells to sites of both acute and chronic inflammation. In this study we demonstrate chronic ibrutinib treatment reduced macrophage accumulation within the diabetic kidney. Using a model of acute inflammatory cell recruitment, we further demonstrate that acute ibrutinib treatment reduced immune cell accumulation via reducing chemokine production, including the chemokine CCL2 which is pivotal for monocyte/macrophage recruitment in many pre-clinical models of inflammation. Both genetic deletion and pharmacological inhibition of CCR2 dramatically reduces macrophage recruitment and protects the kidney from functional decline (49)(50). We show a reduction in macrophage accumulation in the diabetic kidney in mice treated with ibrutinib, and this is coupled with lower expression of CCL2 mRNA, demonstrating that treatment with ibrutinib reduces the production of CCL2 in the diabetic kidney in an NF-κB dependent manner. These results are consistent with previous experimental models of diabetic nephropathy whereby macrophage accumulation is reduced in CCL2 knock out mice.

Having demonstrated that ibrutinib treatment limits immune cell infiltration we went on to show that mice fed a HFD and treated with ibrutinib have a reduction in the activation of both NF-κB and NLRP3 inflammasome in the kidney. Indeed, this was associated with the reduced production of mRNA of the pro-inflammatory cytokines IL-1β, TNFα and IL-18, and the chemokines CXCL1, CCL2 and CCL5. Specific pharmacological inhibition and genetic deletion of NF-κB has been shown to attenuate functional and structural injury in mice in pre-clinical models of acute renal ischemia reperfusion injury (51) and in chronic models of chronic kidney disease (33) and diabetic nephropathy (20)(40). NLRP3/ASC knockout mice consistently display protection against renal injury in both the acute and chronic setting including diabetic nephropathy (52) and western-diet induced diabetic nephropathy (53).

We also demonstrate that treatment of ibrutinib to mice fed a HFD attenuated the formation of the NLRP3 complex, and blocked the proteolytic cleavage of pro-caspase 1 to active caspase 1 in the diabetic kidney and liver. Similarly, caspase 1/11 knock out mice display reduced renal inflammation and improved renal function in models of diabetic nephropathy and lupus nephritis (54). A clinical trial with monoclonal anti–IL-1β IgG gevokizumab in type 2 diabetic kidney disease is ongoing (EudraCT2013-003610-41) suggesting that targeting this pathway could have therapeutic value. Individually targeting the NF-κB and NLRP3 have been shown to be efficacious in protecting organs from functional decline; here we have used an FDA approved medicine, ibrutinib, which targets both NF-κB and NLRP3 inflammasome eliciting protection from microvascular damage.

### Conclusion

The concept that the development of peripheral insulin resistance is intrinsically linked to the extent of myeloid cell activation is emerging in the field of diabetic medicine. Here we demonstrate that by targeting the inflammatory component of diet-induced diabetes, via a novel target BTK, with a FDA approved medicine ibrutinib, we can not only reduce activation of NF-κB and the NLRP3 inflammasome in the diabetic tissue, but dramatically reduce the development of insulin resistance *in vivo*. Ultimately, preserving insulin signalling results in improved glucose homeostasis and protects the diabetic kidney from functional decline. Importantly, we have shown that ibrutinib treatment to murine and human macrophages significantly reduces IL-1β secretion and TNFα secretion by over 90 %. We have identified a FDA approved medication that has many of the ideal properties of candidate medication for repurposing into the treatment of type 2 diabetes and its microvascular complications.

## Methods

### Animals and Human Studies

The experimental protocols used in this study have been approved by the Animal Welfare Ethics Review Board (AWERB) of Queen Mary University of London and the University of Oxford, the study was performed under license issued by the Home Office. Animal care was in accordance with the Home Office guidance on Operation of Animals (Scientific Procedures Act 1986) published by Her Majesty’s Stationery Office.

#### High-fat-diet induced insulin resistance

10 weeks old C57BL/6 mice, housed in the same unit under conventional housing conditions at 25±2 °C were randomly assigned either normal diet (chow) or high-fat-high diet (HFD) (D12331 diet, Research Diet Inc., USA). All mice had access to food and water *ad libitum*. After 6 weeks of dietary manipulation, mice were randomly assigned to a treatment group receiving either ibrutinib (3 or 30 mg/kg, p.o) or vehicle (5 % DMSO, 30 % cyclodextrin, p.o.) for 5 days per week for 6 weeks. The dose used has previously been reported to be selectively inhibit BTK (29). One week prior to the end of the experiment, an oral glucose tolerance test was preformed, 24 h prior to the end of the experiment, mice were placed in metabolic cages and urine collected. Blood was collected by cardiac puncture under general anesthesia.

#### Zymosan induced peritonitis

8-10 weeks old male C57BL/6J mice were pretreated with ibrutinib (10 mg/kg, p.o.) or vehicle (5 % DMSO, 30 % cyclodextrin, p.o.) 1 h prior to administration of 100 µg zymosan A (Sigma-Aldrich) in PBS. After 2, 4, 16 and 48 h mice were sacrificed and peritoneal exudates collected by lavage with 5 ml of ice cold sterile PBS with 2 mM EDTA.

### Cells culture

#### Macrophage reporter cell line (RAW Blue)

A commercially available reporter macrophage cell line (RAW-Blue cells; InvivoGen, San Diego, CA) was used: Briefly, cells were derived from murine RAW 264.7 macrophages with chromosomal integration of a secreted embryonic alkaline phosphatase (SEAP) reporter construct, induced by NF-κB and activator protein 1 (AP-1) transcriptional activation. Cells were grown to 80% confluence in T-25 culture flasks in Dulbecco’s modified Eagle’s medium (DMEM) containing 4.5 g/liter glucose, heat-inactivated 10% fetal bovine serum, 2 mM L-glutamine, and 200 µg/ml Zeocin antibiotic (InvivoGen) at 37°C in 5% CO_2_. To minimize experimental variability, only cells with fewer than 5 passages were used. Cells were plated at 0.95×10^5^ per well. After experimental treatment 20 ul of cell supernants was added to 180 ul of QuantiBlue substrate (InvivoGen) and incubated at 30 C for 60 mins, plate was then read at an optical density at 655 nm (OD_655_) on a microplate spectrophotometer (PherastarFSX, BMG Lab, UK).

#### Murine Bone Marrow-Derived Macrophages (BMDMs)

Bone marrow-derived macrophages were generated as previously described (55). Briefly, fresh bone marrow cells from tibiae and femurs of male C57BL/6 mice aged 8-10 weeks were cultured in Dulbecco’s Modified Eagle’s medium (DMEM) containing 4.5 g/liter glucose, 2 mM L-glutamine, 50 units/ml penicillin and 50 µg /ml streptomycin, 10% heat-inactivated fetal bovine serum (FBS), 10% L929 cell-conditioned media (as a source of macrophage colony-stimulating factor) and for 7 days. Bone marrow cells were seeded into 8 ml of medium in 90 mm non-tissue culture treated Petri dishes (ThermoFisher Scientific, Sterilin, UK). On day 5, and additional 5 ml of medium was added. Gentle scrapping was used to lift cells. BMDMs were then counted and suspended in FBS free media at the desired cell concentration.

#### Human monocyte-derived macrophages (hMoDMs)

Peripheral blood mononuclear cells (PBMC) were purified from leukocyte cones from healthy volunteers with informed consent (NHSBTS, Oxford, UK) by density centrifugation over Ficoll-Paque PLUS (Sigma). The PBMC layer was carefully harvested and then washed twice with PBS. Monocytes were then isolated from the PBMCs by negative selection using magnetic beads (Miltenyi Biotec, Bergisch Gladbach, Germany). Cells were maintained in RPMI 1640 medium supplemented with 1 % human serum, 50 ng/ml macrophage colony stimulating factor (hM-CSF, BioLegend, San Diego, USA), 50 units/ml penicillin and 50 µg /ml streptomycin for 7 days.

### Caspase 1 assay

Caspase 1 activity was measured using commercially available kit (Caspase 1 Glo inflammasome assay, Promega). Briefly, BMDM (2.5 x10^4^) were seeded on half area white walled 96 well multiwall plates. BMDM where pre-treated with ibrutinib for one hour prior to 8 h LPS (100 ng ml^-1^) stimulation. For caspase 1 activity add 5 mM ATP in the last 60 min then incubate 20 ul cell supernant with 40µM Z-WEHD-aminoluciferin 120µM MG-132 inhibtor in Caspase-Glo® 1 Buffer for 1 h then read luminescence using microplate spectrophotometer (PherastarFSX, BMG Lab).

### mRNA expression

Mouse tissues and cultured cells were harvested into TRIzol reagent (Life Technologies) and total RNA extracted. RNA concentration and quality was determined with a ND-1000 spectrophotometer (Nano Drop Technologies, Wilmington, USA). cDNA was synthesized from 1000 ng RNA using the QuantiTect Reverse Transcription kit (Qiagen, Manchester, UK) according to the manufacturer’s instructions. Real-time quantitative PCR was performed using Sybr Select gene expression master mix (Life Technologies) in the StepOnePlus^TM^ thermal cycler (Applied Biosystems). Primers were purchased from Qiagen (*il1β, il18, tnfα, il6, il10, cxcl1*, *ccl2, ccl5 18S*, *βactin*). Cycle threshold values were determined by the StepOne software and target gene expression was normalized to housekeeping gene (*βactin or 18S*).

### Quantification of cytokine level

BMDM or hMoDM (1.5×10^6^) were seeded in 6 well plated. BMDM or hMoDM were pre-treated with ibrutinib for one hour prior to 8 h LPS (100 ng ml^-1^) stimulation. Culture medium was adjusted to 5 mM in the last 60 min to allow secretion of IL-1β.

*ELISA:* measurement of IL-1β, TNFα and IL-6, CXCL1, CCL2 and CCL5 secreted protein levels in cell supernatants from 1×10^6^ murine BMDM’s was performed by ELISA (R&D Systems) according to manufactureŕs instructions.

*Bead assay*: measurement of IL-1β, TNFα, and IL-6 secreted protein level in cell supernatant from 1×10^6^ human MoDM’s was performed by Legend Plex multi-analyte assay kit (BioLegend) according to manufacturer’s instruction. Data were acquired using a BD Fortessa X20 cytometer (BD Biosciences), and analyzed using LegendPlex Software (BioLegened).

### Flow cytometry

Cells were washed in FACS buffer (0.05 % BSA, 2 mM EDTA in PBS pH 7.4) blocked using anti CD16/32 for 10 mins at 4°C, followed by antibody staining for the surface markers BV421-conjugated CD11b (BioLegend), BV510-conjugated Ly6G (BioLegend), PeCP-conjugated LyGC (BioLegend), PE-conjugated CD14^+^ (BioLegend) with appropriate isotype controls. Absolute cell counts were performed using quantity of calibration beads added to each sample (CountBright, Invitrogen). Data were acquired using a BD Fortessa X20 cytometer (BD Biosciences) and then analysed using FlowJo (Tree Star Inc, USA) software.

### Western Blot

BMDM or tissue (kidney and liver) were lysed in presence of protease and phosphatase inhibitors (Sigma-Aldrich) spun (14000 g for 15 min) and supernant collected as previously described (56). Protein concentration was determined by using a BCA protein assay kit (ThermoFisher Scientific). Total cell protein (30 mg) was added to 4x Laemmli buffer (250 mM Tris-HCl, pH 6.8, 8% SDS, 40% glycerol, 0.004% bromophenol blue, 20% β-mercaptoethanol) and heated at 95°C for 5 min. Samples were then resolved on SDS-PAGE gels and transferred onto Hybond ECL nitrocellulose membranes (GE Healthcare, Buckinghamshire, UK). Membranes were blocked with 5% milk in TBS-T for 1 hour at RT and then incubated with the primary antibody (rabbit anti-Ser^473^-Akt, anti-total Akt, mouse anti-Thr^202^/Tyr^204^-ERK1/2, rabbit anti-total ERK1/2, rabbit, rabbit anti-Thr^183^/Tyr^185^-JNK, rabbit anti-total JNK, rabbit anti-Ser^307^-IRS1, mouse anti-total IRS1, rabbit anti-phospho-Ser^9^-GSK-3β, rabbit anti-total GSK-3β, mouse anti-Ser^32/36^-IKBα, mouse anti-total IKBα, rabbit anti-Ser^176/180^-IKKα/β, rabbit anti-total IKKα/β, rabbit anti-NF-κB, mouse anti-caspase 1 (p20)) diluted 1:1000 in 2% BSA/TBS-T overnight at 4°C. Next, membranes were incubated with an HRP-conjugated anti-rabbit or anti mouse secondary antibody diluted 1:10000 in 2 % BSA for 1 hour at RT. Protein bands were visualized by incubating the membranes for 5 min with Amersham ECL prime and subsequent exposure to X-ray film over a range of exposure times. β-actin and histone 3 were used as loading control. For successive antibody incubations using the same membrane bound antibodies were removed with stripping buffer (ThermoFisher Scientific).

### Histology

*Immunofluorescence:* for p-65 expression and location detection by immunofluorescence, cells placed in a 4-well chamber-slide were fixed (4% paraformaldehyde) for 10 minutes at RT, permeabilized (0.5% Triton X-100 in PBS) for 10 minutes at 4°C, blocked with 4% BSA/PBS containing 6% donkey serum, and incubated with mouse anti-p-65 antibody diluted 1:100 in 4% BSA/PBS, overnight at 4°C. Cells were then incubated with donkey anti-mouse Alexa Fluor 488 diluted 1:500 in 4% BSA/PBS for 1 hour at RT. DAPI (4’,6-diamidino-2-phenylindole) was used for nuclear counterstaining. The slide was mounted with Fluormount-G® and cells were visualized using a confocal microscope.

*Oil Red O staining:* Frozen liver samples were embedded in OCT, and cut in 10 µm sections. Section were brought to room temperature, fixed with 10 % buffer formalin for 5 min, washed with 60 % isopropanol, then saturated with Oil Red O (1% w/v, 60 % isopropanol) for 15 min, washed in 60 % isopropanol and rinsed in distilled water. Then sections were then mounted in aqueous mounting medium with coverslips. Images were acquired using a NanoZoomer Digital Pathology Scanner (Hamamatsu Photonics K.K., Japan) and analyzed using the NDP Viewer software.

*Periodic Acids Schiff’s staining:* Kidney samples were obtained at the end of the experiment and fixed in 10 % neutral-buffered formalin for 48 h and histology staining was performed. Briefly, kidney tissue was embedding in paraffin and processed to obtain 4 µm sections. After deparaffinization and sections were rehydrated through graded alcohol to distilled water. The sections were then incubated with saturated in Periodic Acid Schiff’s (Sigma, UK) solution for 30 min and washed in distilled water. Then sections were then dehydrated through graded alcohols and cleared before mounting with coverslips. Images were acquired using a NanoZoomer Digital Pathology Scanner and analyzed using the NDP Viewer software.

*Immunohistochemistry:* Kidney sections cut at 4 µm were deparaffinized to PBS. Antigen retrieval was performed by in citrate buffer (pH 6.0) for 15 minutes. Once cooled, sections were incubated with 3% H_2_O_2_ for 20 minutes to inactivate endogenous peroxidases (Dako EnVision+ System-HRP-DAB, K4010) and subsequently treated with 10% normal goat serum (Dako, UK) to reduce nonspecific absorption. Sections were subsequently incubated at 37°C for 1 hour with the following primary antibodies: anti-F4/80 (1:400, cat#ab5694, Abcam, UK) washed with PBS, and then incubated at room temperature for 30 minutes with labelled polymer-HRP antibody (Dako EnVision+ System, HRP-DAB). Sections were developed in DAB chromogen solution, and the reaction stopped by immersion of sections in water. Counterstaining was performed with Harris hematoxylin before sections were dehydrated and mounted in DPX mounting medium.

### Statistical analysis

All data in the text and figures are presented as mean ± standard error mean (SEM) of n observations, where n represents the number of animals studied (*in vivo*) or biological replicates (*in vitro*). All statistical analysis was calculated using GraphPad Prism 7 for Mac (GraphPad Software, San Diego, California, USA). Data without repeated measurements was assessed by a one-way ANOVA followed by Bonferroni correction. In all cases a p<0.05 was deemed significant.

### Data and Resource Availability

The datasets generated during and/or analyzed during the current study are available from the corresponding author upon reasonable request. No novel resources were generated during the current study.

### Author contribution statement

G.S.D.P., M.C., H.M.A.T., F.C., C.E.O. and L.Z. performed experiments; G.S.D.P., M.C. and H.M.A.T. analyzed results and made the figures; G.S.D.P., P.B., N.G., D.R.G. and C.T., designed the research. G.S.D.P, D.R.G. and C.T. wrote the paper. All authors provided critical revision of the manuscript.

## Acknowledgements

We would like to thank the following funding bodies for their support: the British Heart Foundation (Award number: FS/13/58/30648) to G.S.D.P., (Award number: RG/15/10/23915) to D.R.G and the BHF Centre of Research Excellence (RE/13/1/30181) to G.S.D.P and D.R.G. Queen Mary University of London to C.E.O. William Harvey Research Foundation to CT; Bart’s and The London Charity Centre of Diabetic Kidney Disease (program grant: 577/2348) to CT.

