## Supplementary data for "Inhibition of Bruton’s tyrosine kinase reduces NF-kB and NLRP3 inflammasome activity preventing insulin resistance and microvascular disease"

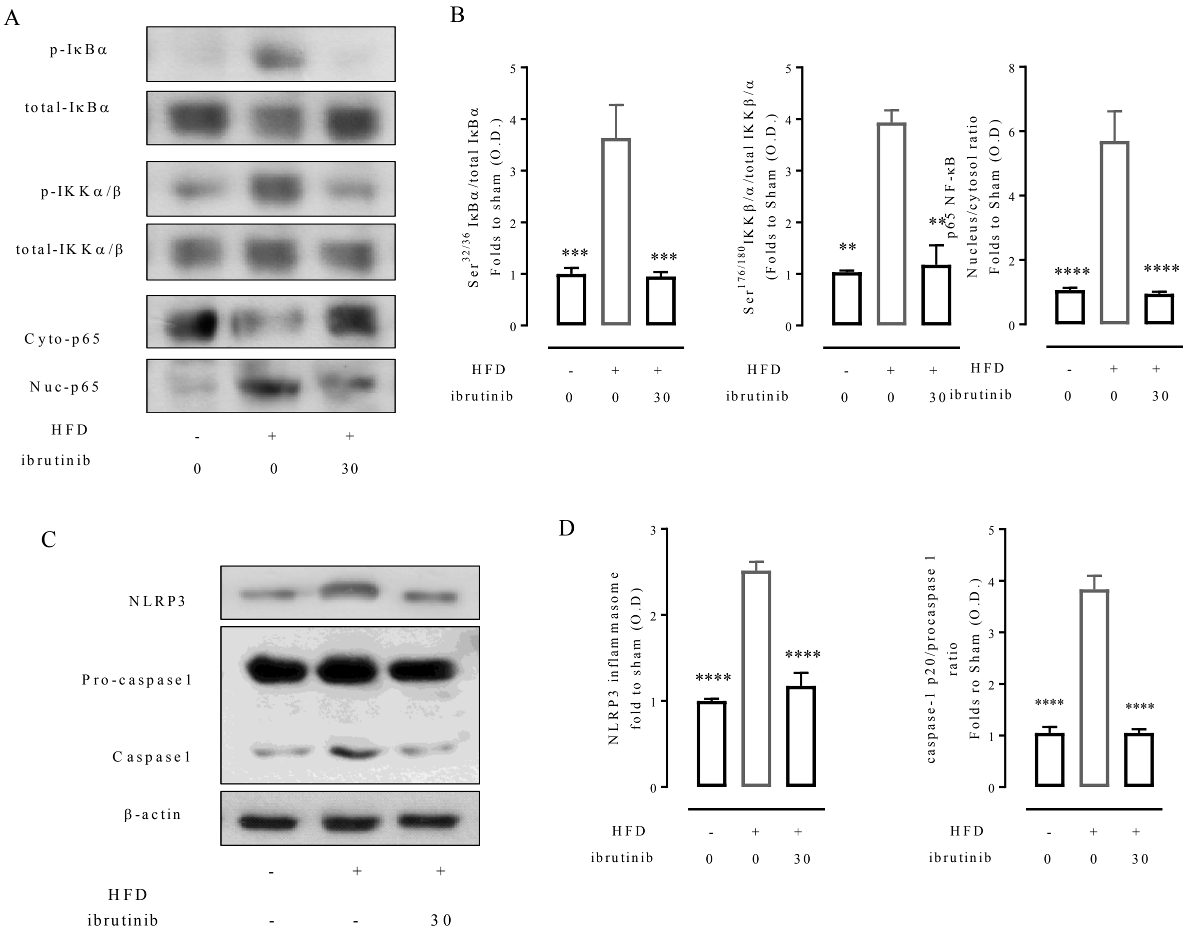


**Supplementary figure 1: Ibrutinib treatment reduces inflammation in the diabetic kidney via inhibition of NF-κB and the NLRP3 inflammasome.**

C57BL/6 mice, fed a standard diet (chow) or a high-fat diet (HFD) for 12 weeks, were treated with vehicle or ibrutinib (3 o 30 mg/kg) five times per week between weeks 7 and 12.A) Representative western blots for phosphorylation of $Ser^{32/36}$ on IKBα in the kidney and normalized to total IKBα; for phosphorylation of $Ser^{176/180}$ on IKKα in the kidney and normalized to total IKKα; nuclear translocation of p65 and B) quantified using densitometry. C) Representative western blots for NLRP3 inflammasome assembly and the proteolytic cleavage of pro-caspase 1 to caspase 1 normalized to β-actin and D) quantified using densitometry. Data analyses by a one-way ANOVA followed by a Bonferroni *post-hoc* test and the mean is expressed mean± SEM of n=9/10 per group. **p< 0.01, ***p<0.001 and ****p<0.0001.

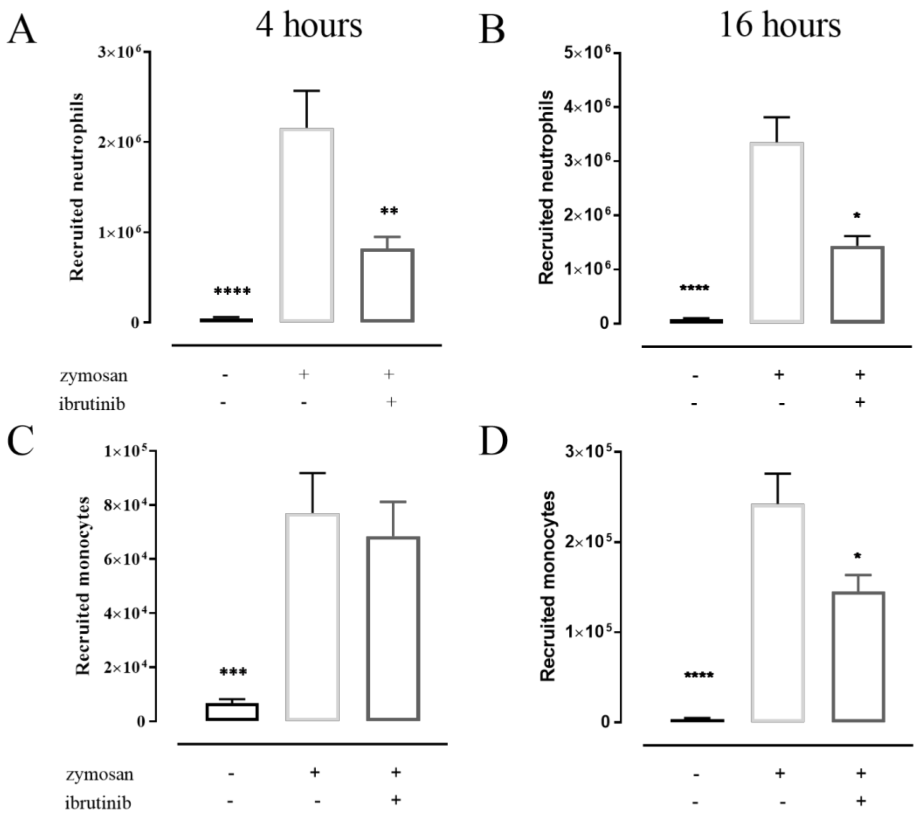


**Supplementary figure 2:**

C57BL/6 mice were pre-treated with ibrutinib (10 mg/kg, p.o.) or vehicle 1 h prior to zymosan challenge (100 µg, i.p.). Mice were euthanized at 4, 16 and peritoneal lavage samples were collected. Quantification of recruited neutrophils at A) 4 hour and B) 16 hours; and monocytes at ) 4 hour and D) 16 hours post zymosan injection. Data shown are mean± SEM of n=12 per group *p< 0.05 **p< 0.01.

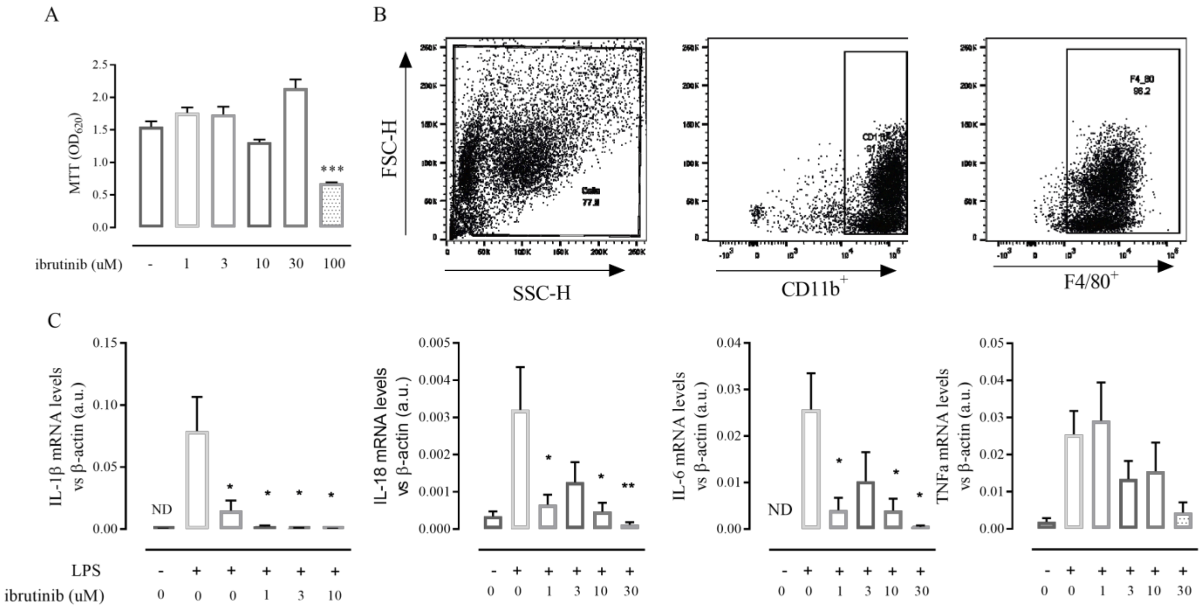


**Supplementary figure 3:**

A) Assessment of cytotoxicity of ibrutinib with MTT assay in RAW blue cells. B**)** Representative flow cytometry plots of BMDM differentiation at day 7. C) Relative gene expression of *IL-1β, IL-18, IL-6 and TNFa* were assessed by qPCR and normalized to 18S. *p<0.05,**p< 0.01, ***p<0.001.
